# Assessment of Q-PCR based vector copy numbers and viral titration in CAR T cell products, Probe vs SYBR Green

**DOI:** 10.64898/2026.10.19.760184

**Authors:** Hamidreza Ebrahimiyan, Ali Sayadmanesh, Azadeh Moradmand, Hossein Baharvand, Mohsen Basiri

**Affiliations:** Department of Stem Cells and Developmental Biology, Cell Science Research Center, Royan Institute for Stem Cell Biology and Technology, ACECR, Tehran, Iran; Department of Molecular Medicine, Faculty of Advanced Technologies in Medicine, Iran University of Medical Sciences, Tehran, Iran; Advanced Therapy Medicinal Product Technology Development Center (ATMP-TDC), Royan Institute for Stem Cell Biology and Technology, ACECR, Tehran, Iran; Department of Developmental Biology, School of Basic Sciences and Advanced Technologies in Biology, University of Science and Culture, Tehran, Iran

**Keywords:** Chimeric Antigen Receptor (CAR), Genetic engineering, T cell therapy, Vector Copy Numbers (VCNs), Viral titration, Q-PCR, Vector Bank (VB)

## Abstract

Assessment of vector copy numbers (VCNs) in comprised of the chimeric antigen receptor (CAR) T cells as a safety parameter and viral titration in the vector bank (VB) as an efficacy parameter are essential in the CAR T cell production procedure. Here, we developed a quantitative-PCR (Q-PCR) based VCN and viral titration analysis using both Probe and SYBR Green methods. Cellular viral titration was also applied for comparison of viral titration results. The VCN in CAR T cells with SYBR Green was 1.72, 2.31, and 2.46 respectively, while in the Probe based analysis, it was 1.33, 1.83, and 2.20. The molecular viral titration in SYBR Green resulted in 2.90×10^9^, 5.24×10^10^, and 9.76×10^10^, while in the Probe it was 1.54×10^9^, 1.13×10^9^, and 7.73×10^8^. The cellular titration resulted in 1.73×10^6^, 2.16×10^6^, and 2.49×10^6^.

In summary, both SYBR Green and Probe revealing that Probe strategy had a lower dependency on sample contamination and could be a better strategy for the Q-PCR method. The SYBR Green based viral titration method is significantly different from the Probe based method, with the Probe based method yielding better results compared to the cellular titration method. However, no significant difference was reported in both methods for VCNs.

## Introduction

Immunotherapy is an advanced treatment that enhances the body’s natural defense against cancer. Adoptive cell therapy (ACT) has garnered increased interest from oncology and immunology experts in the past decade. ACT is a personalized immune therapy in which the patient’s own immune cells are enriched or genetically modified, expanded in vitro, and reinfused to combat tumors. Chimeric antigen receptor (CAR) T cells have made significant progress in ACT for malignant diseases (1). Chimeric Antigen Receptors (CARs) are engineered molecular constructs that incorporate a designated antibody-derived single-chain variable fragment (scFv) functioning as an extracellular domain to facilitate the recognition of specific antigens (2). The scFv is linked to various intracellular domains that initiate intracellular signaling in CAR T cells, crucial for activation, proliferation, and cytotoxicity (3). Different CD19-targeted CAR T cell therapies have demonstrated promising clinical outcomes in lymphoid malignancies such as chronic lymphocytic leukemia (CLL), acute lymphoblastic leukemia (ALL), and non-Hodgkin’s lymphoma (NHL) (4). Additionally, the application of CAR T cell therapies is being investigated in the context of solid tumors and other lymphoid disorders (5). A range of anti-CD19 CAR T cell therapeutic strategies have received authorization from the United States Food and Drug Administration (FDA) as well as the European Medicines Agency (EMA). The production of CAR T cells involves introducing cDNA encoding the CAR receptor into the T cells. Most strategies used to insert genes into the genome for long-term expression have resulted in increased expansion of gene-modified cells both in vitro and in vivo (6). Recombinant retrovirus and lentivirus are commonly used in CAR T cell therapies, classified as gene therapy medicinal products (GTMP) due to the genetic modification of CAR T cells. CAR T cell products are manufactured according to good manufacturing practice (GMP) standards (6). Extensive safety evaluations and quality control tests are necessary for advanced therapy medicinal products (ATMP) like CAR T cell therapy.

Retroviral vectors are randomly inserted into the genome and require quality control for the safety of CAR T cell products (7). The number of transgene copies in CAR T cells (vector copy number, VCN) must be assessed before injection to patients to prevent genotoxicity risks (8). The assessment of the proliferation and longevity of CAR T cells within patients is essential for the comprehensive evaluation of therapeutic responses to CAR T cell interventions. Quantitative PCR (Q-PCR) is typically used to assess VCN in CAR T cell products as a quality control test (9–12). A VCN of <5 is recommended as safe for treatment in clinical applications (13). For clinical application, retrovirus production must be done under GMP conditions, with potency characterized by viral titration (7). Each CAR T cell product with an acceptable VCN should report the viral multiplicity of infection (MOI) to prevent genotoxicity risks. The MOI is the number of viral particles that can infect each cell, which a higher MOI potentially increasing VCN and genotoxicity risks (14). Determining the exact MOI and VCN, along with related genotoxicity tests, is essential for each CAR T cell product. Viral titration and MOI can be determined through cellular-based and molecular-based methods to ensure accurate results (15).

We developed a Q-PCR method using both the current SYBR Green and Probe based absolute standard curve method (ACM) to determine VCN and MOI. The primers targeted the U5 region in 5’ LTR, applicable not only for CAR but also on the same SFG vector backbone. Using the same primers, standards, and Q-PCR conditions reduced time and costs for VCN and MOI experiments. Both the SYBR Green and Probe methods were analyzed and compared to optimize experiments. For MOI analysis, both molecular and cellular tests were applied and compared for precision.

## Methods and materials

### Vector generation

The CAR plasmid was sourced from the Royan Plasmid Bank. The TOP10 E. coli cells were transformed and cultured for CAR plasmid extraction. The Plasmid Maxi Extraction Kit (QIAGEN, cat: 12123) was used for the preparation of the CAR plasmid.

### Cell Lines, Culture Conditions, and T Cell Isolation

The Plat-A (Platinum-A) and PG13 (ATCC CRL-10686) cell lines were cultured in IMDM (Iscove’s Modified Dulbecco’s Medium, Gibco cat: 21980032) with 10% HyClone Characterized Fetal Bovine Serum (FBS, Cytiva cat: SH30071.03IH30-45) and 2 mM GlutaMax (100x, Gibco cat: 35050-061) at 37°C in 5% CO2. Peripheral Blood Mononuclear Cells (PBMCs) from three healthy donors were separated by centrifugation on Lymphoprep (Serumwerk Bernburg cat: 1114544). On day 0, PBMCs were activated on an anti-CD3 and anti-CD28 (Miltenyibiotec, cat: 130-093-387 and 130-093-375) pre-coated 24-well plate in CTL media containing 45% RPMI-1640 (Cytiva cat: SH3009601), 45% Click’s medium (Irvine Scientific cat: 9195), 10% FBS, and 2 mM GlutaMax. On day 1, interleukin-2 (IL-2, R&D Systems cat: 202-IL-010/CF) was added to the culture at a final concentration of 100 IU/mL and repeated every two days to maintain the culture.

### Transfection, Virus Production, and T Cell Transduction

To transiently produce retroviruses, Plat-A cells were transfected with a CAR plasmid using Lipofectamine 3000 (Invitrogen, Part No. 100022234) following the manufacturer’s instructions. The transfected cells were incubated overnight at 37°C. The retroviral supernatant was harvested 24 hours post-replacement, subsequently filtered through a 0.45 mm filter, snap-frozen utilizing dry ice, and preserved at a temperature of −80° For PG13 transduction. On day 0, 4×10^4^ cells were seeded in a 24-well plate. On day 1, the media was replaced by retroviral supernatant with 10 μg Polybrene (MERK, SKU: TR-1003-G). After overnight incubation, the cell media was replaced with fresh media. When cell confluences reached 80-90%, the transduction procedure was repeated until 80% of PG13 cells were CAR^+^. The CAR PG13 cells were sorted, and a positive single clone was isolated to use as a producer cell line. To produce retroviral particles, the CAR PG13 cells were seeded at a density of 43 × 10^3^/cm^2^. After 48 hours, the fresh medium was replaced, and 24 hours later, the supernatant containing retroviral particles was collected, filtered, snap-frozen on dry ice, and stored at −80°C for T cell transduction. To produce CAR T cells, on day 3, after T cell activation, 2.5 mL of retroviral supernatant was transferred to a non-treated 24-well plate pre-coated with Retronectin (Takara cat: T100A/B) and centrifuged for 2 hours at 2000 × g at 4°C. Following centrifugation, the supernatant was removed, and the activated T cells (2 × 10^5^ cells) were re-suspended in a complete medium supplemented with IL-2 (100 IU/mL), introduced into the wells, and subsequently centrifuged at 400 × g for a duration of 10 minutes. On day 5, CAR T cells were harvested for the experiment.

### DNA, Viral RNA Extraction, and Standard Production

The QIAamp DNA Mini Kit (QIAGEN, cat: 51301) was used to extract cellular DNA according to the manufacturer’s protocol for use in conventional PCR. The QIAamp Viral RNA Mini Kit (QIAGEN, cat: 52904) and QuantiTect Reverse Transcription Kit (QIAGEN, cat: 205311) were used to extract viral RNA and synthesize the cDNA for use in Q-PCR. The DNA extracted from CAR T cells was used to produce standards for copy number detection. Conventional PCR was conducted using 10 μl Taq DNA Polymerase 2x Master Mix RED (Cat no. A190307, Ampliqon), 2 μl DNA template, 1 μl each two-set primer (Table 1), and ribonuclease-free water to a total volume of 20 μl. The conditions of PCR were as follows: 95°C for 15 min, 35 cycles of 95°C for 15 seconds, 56°C for 30 seconds, 72°C for 1 min, and then 72°C for 15 min. PCR products were run on a 0.5% agarose gel and recovered using the QIAquick Gel Extraction Kit. The concentration of the extracted DNA was measured three times with the Qubit dsDNA Quantification Assay Kit (Invitrogen, code: 10616763) to calculate the absolute copy number (Table 2).

**Table 1:**
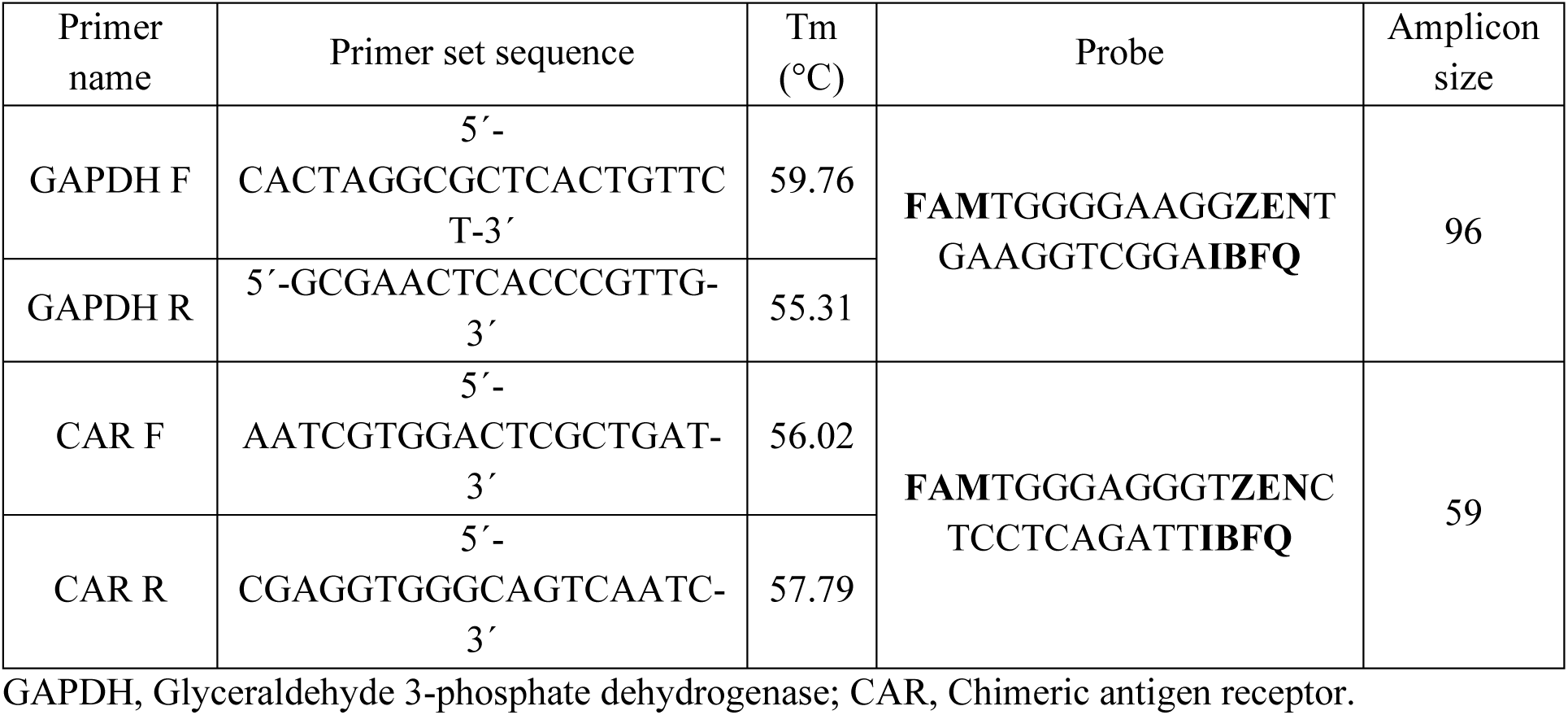
Details of primers.

**Table 2:** Copy number SYBR Green and Probe (sample1)

| Initial sample | Dilution factor | Standard concentration average (ng/μl) | Absolute copy number (copies/μl) | Q-PCR Ct GAPDH average | Q-PCR Ct CAR average | Detected copy number (copies/μl)<br>*<br>GAPDH | Detected copy number (copies/μl)<br>*<br>CAR | Copy number ratio** (CAR<sup>+</sup> cells) |
|---|---|---|---|---|---|---|---|---|
| SYBR Green |  |  |  |  |  |  |  |  |
| Standard | 10<sup>-2</sup> | 1×10<sup>-2</sup> | 8.69×10<sup>7</sup> | 15.17 | NA | 8.86×10<sup>7</sup> | NA | NA |
| Standard | 10<sup>-3</sup> | 1×10<sup>-3</sup> | 8.69×10<sup>6</sup> | 18.67 | NA | 8.57×10<sup>6</sup> | NA | NA |
| Standard | 10<sup>-4</sup> | 1×10<sup>-4</sup> | 8.69×10<sup>5</sup> | 22.15 | NA | 8.40×10<sup>5</sup> | NA | NA |
| Standard | 10<sup>-5</sup> | 1×10<sup>-5</sup> | 8.69×10<sup>4</sup> | 25.51 | NA | 8.92×10<sup>4</sup> | NA | NA |
| Standard | 10<sup>-2</sup> | 1.8×10<sup>-2</sup> | 2.83×10<sup>8</sup> | NA | 12.96 | NA | 2.93×10<sup>8</sup> | NA |
| Standard | 10<sup>-3</sup> | 1.8×10<sup>-3</sup> | 2.83×10<sup>7</sup> | NA | 16.50 | NA | 2.76×10<sup>7</sup> | NA |
| Standard | 10<sup>-4</sup> | 1.8×10<sup>-4</sup> | 2.83×10<sup>6</sup> | NA | 19.99 | NA | 2.66×10<sup>6</sup> | NA |
| Standard | 10<sup>-5</sup> | 1.8×10<sup>-5</sup> | 2.83×10<sup>5</sup> | NA | 23.27 | NA | 2.97×10<sup>5</sup> | NA |
| CAR T cell | <sup>1</sup>/<sub>2</sub> | NA | NA | 21.53 | 20.55 | 1.27×10<sup>6</sup> | 1.84×10<sup>6</sup> | 1.44 (1.72) |
| CAR T cell | <sup>1</sup>/<sub>10</sub> | NA | NA | 23.33 | 22.39 | 3.82×10<sup>5</sup> | 5.36×10<sup>5</sup> | 1.40 (1.67) |
| Probe |  |  |  |  |  |  |  |  |
| Standard | 10<sup>-2</sup> | 1×10<sup>-2</sup> | 8.69×10<sup>7</sup> | 17.93 | NA | 8.92×10<sup>7</sup> | NA | NA |
| Standard | 10<sup>-3</sup> | 1×10<sup>-3</sup> | 8.69×10<sup>6</sup> | 21.40 | NA | 8.61×10<sup>6</sup> | NA | NA |
| Standard | 10<sup>-4</sup> | 1×10<sup>-4</sup> | 8.69×10<sup>5</sup> | 24.89 | NA | 8.15×10<sup>5</sup> | NA | NA |
| Standard | 10<sup>-5</sup> | 1×10<sup>-5</sup> | 8.69×10<sup>4</sup> | 28.14 | NA | 9.09×10<sup>4</sup> | NA | NA |
| Standard | 10<sup>-2</sup> | 1.8×10<sup>-2</sup> | 2.83×10<sup>8</sup> | NA | 15.15 | NA | 2.85×10<sup>8</sup> | NA |
| Standard | 10<sup>-3</sup> | 1.8×10<sup>-3</sup> | 2.83×10<sup>7</sup> | NA | 18.64 | NA | 2.74×10<sup>7</sup> | NA |
| Standard | 10<sup>-4</sup> | 1.8×10<sup>-4</sup> | 2.83×10<sup>6</sup> | NA | 21.98 | NA | 2.92×10<sup>6</sup> | NA |
| Standard | 10<sup>-5</sup> | 1.8×10<sup>-5</sup> | 2.83×10<sup>5</sup> | NA | 25.49 | NA | 2.79×10<sup>5</sup> | NA |
| CAR T cell | <sup>1</sup>/<sub>2</sub> | NA | NA | 23.89 | 22.73 | 1.60×10<sup>6</sup> | 1.77×10<sup>6</sup> | 1.11 (1.33) |
| CAR T cell | <sup>1</sup>/<sub>10</sub> | NA | NA | 26.24 | 25.10 | 3.28×10<sup>5</sup> | 3.61×10<sup>5</sup> | 1.10 (1.32) |

NA, not applicated; CAR, Chimeric antigen receptor; \*: The calculated copy number; \*\* The mean copy number of CAR in each cell CAR/GAPDH in total and CAR<sup>+</sup> cells.

### Copy Number Detection and Molecular Viral Titration (Q-PCR)

The GAPDH and CAR standards were diluted with serial dilution (10^−2^ to 10^−6^). The CAR T cell DNA and viral cDNA were also diluted in this procedure. The GAPDH standard alongside CAR T cell DNA were amplified utilizing the GAPDH primer, while the CAR standard and CAR T cell DNA underwent amplification with the CAR primer to detect the copy number of CAR present within the CAR T cell DNA. The CAR standards and viral cDNA were amplified with the CAR primer to detect the copy number of the virus. The detected copy numbers were calculated based on the standard curve linear formula. Two different procedures were run in Q-PCR, including Probe based and SYBR Green-based, to compare the copy number and viral titer results. The conditions of Q-PCR were as follows: 95°C for 15 min, 45 cycles of 95°C for 15 seconds, 56°C for 30 seconds, and 72°C for 1 min. Q-PCR was performed with 10 μl RealQ Plus 2x Master Mix Green (Ampliqon, cat no. A325402) or RealQ Plus 2x Master Mix for Probe (Ampliqon, cat no. A315402), 2 μl cDNA or DNA template, 1 μl each two-set primer (1 μl Probe) Table 1, and ribonuclease-free water to a total volume of 20 μl. To detecte the copy number of CAR within the CAR T cell, the detected copy number (copies/μl) of CAR was divided by the detected copy number of GAPDH present in the CAR T cell. The original concentration of viruses was also calculated based on the described formula (8).

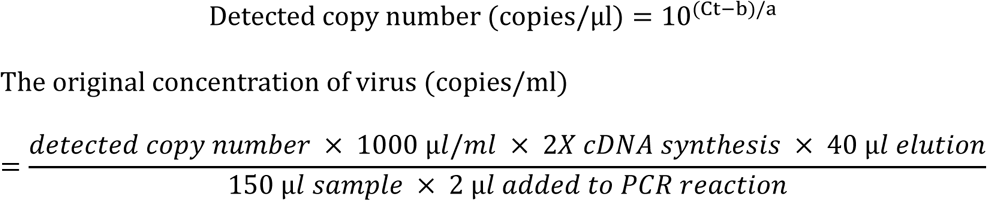

### Cellular viral titration and flow cytometry analysis

To analyze functional viral titers, the original virus sample was diluted at 1/20 and 1/40, and then used for transduction of T cells. For T cell transduction, on day 3, 0.1 mL of retroviral supernatant was added to a non-treated 96-well plate pre-coated with Retronectin and centrifuged for 2 hours at 2000×g at 4°C. The activated T cells (8×10^4^ cells) were resuspended in complete media with IL-2 (100 IU/mL) and then added to the wells, followed by centrifugation at 400×g for 10 minutes.

For analysis of CAR expression, on day 5, cells were collected, washed, and stained with the recombinant CD19 protein FITC (Acro, cat: CD9-HF251), the anti-human CD3 antibody APC (BioLegend, cat: 300311), and their respective isotypes (Acro, cat: DNP-FM2 and BioLegend, cat: 400219) at 4°C for 45 minutes in the dark. All samples were analyzed using BD FACSCalibur flow cytometers (BD Biosciences, Franklin Lakes, NJ, USA). Signals between 1% to 20% were accepted for viral titration calculations (16).

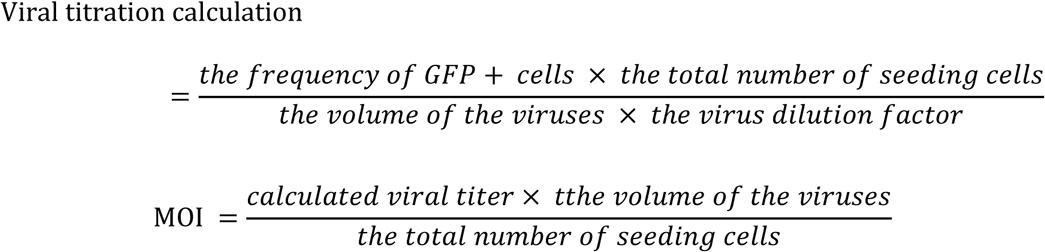

### Statistical analysis

The GraphPad Prism version 10 software (GraphPad Software, La Jolla, CA, www.graphpad.com) was used for plotting and analyzing. Flow cytometry data were analyzed with FlowJo software (Tree Star Inc., Ashland, USA). Linear regression tests were applied, with P < 0.05 and R_2_ > 0.98 defined as a statistically significant level. The unpaired t-test was applied for the comparison of two different methods.

## Results

### Molecular copy number detection

Primers were designed on the U5 region, present in proviral DNA for CAR T cell copy number calculation and viral RNA for molecular viral titration (Figure 1A-C).

**FIGURE 1:**
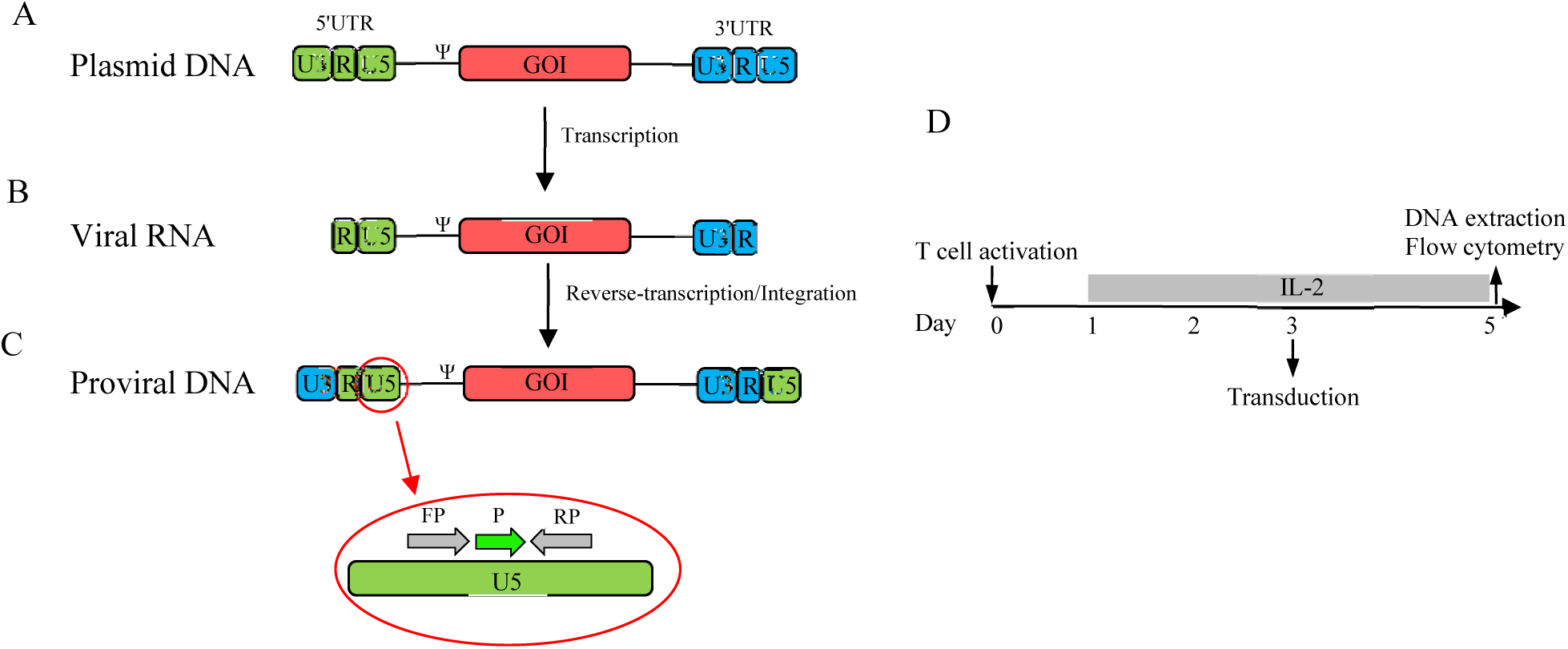
Genome structure of MSCV/MLV based vector and design principle of primers. **Panel A.** Typical plasmid structure of MSCV/MLV retroviral virus production. The 5′ untranslated region (UTR) containing the redundant (R), the unique 5′ (U5) and the unique the 3′ (U3) region, the 3′UTR containing the unique 3′ (U3), unique 5′ (U5) and R region. The ψ (packaging signal). **Panel B.** The structure of retroviral MSCV/MLV viral RNA. **Panel C.** The structure of proviral MSCV/MLV DNA. The forward primer (FP), reverse primer (RP) and probe (P) were designed in U5 region which could be used for viral titration and copy number detection. The proviral DNA have two U5 region however the viral RNA has one copy. **Panel D.** The schematic timeline of the experimental setting. PBMC was activated by anti-CD3 and anti-CD28 dynabead on day 0. The CAR construct was transduced into T cells on the day 3. Flow analysis and DNA extraction applied on day 5.

### SYBR Green method

After CAR T cell DNA extraction on day 5, Q-PCR was conducted (Figure 1D). The amplification plots for GAPDH and CAR standards are shown in Figure 2A and D. A strong linear regression with R_2_ = 0.9999, PCR efficiency of 94.91%, and regression *Pv* < 0.0001 were shown in the GAPDH standard curve, demonstrating a relationship between the Ct values and DNA absolute copy numbers (log scale) in serial dilution (Figure 2C). The detected copy number (copies/ul) in the GAPDH standard = 10^(Ct-42.589)/-3.45^. The detected copy number of GAPDH in a 1∕2 dilution of CAR T cells was calculated at 1.27×10^6^ and for a 1∕10 dilution was 3.82×10^5^. In the CAR standard, a strong linear regression with R_2_ = 0.9997, PCR efficiency of 95.22%, and a regression *Pv* of 0.0002 were shown in the standard curve for serial dilution (Figure 2F). The detected copy number in the CAR standard = 10^(Ct-42.113)/-3.442^. The detected copy number in the 1∕2 dilution of CAR T cells was calculated at 1.84×10^6^ and for 1∕10 dilutions was 5.36×10^5^. The copy number ratio of CAR T cells in 1∕2 and 1∕10 were 1.44 and 1.40 in total cells and 1.72 and 1.67 in CAR^+^ cells (83.60%), respectively (Table 2 and Figure 6 A-B). The amplification plot of CAR T cell samples alongside the first and last standard dilutions is shown in Figure 3 A-B. For copy number calculation based on SYBR Green method, two different dilutions of samples revealed approximately the same result. The detected copy number for CAR and GAPDH standards also revealed precise calculations.

**FIGURE 2:**
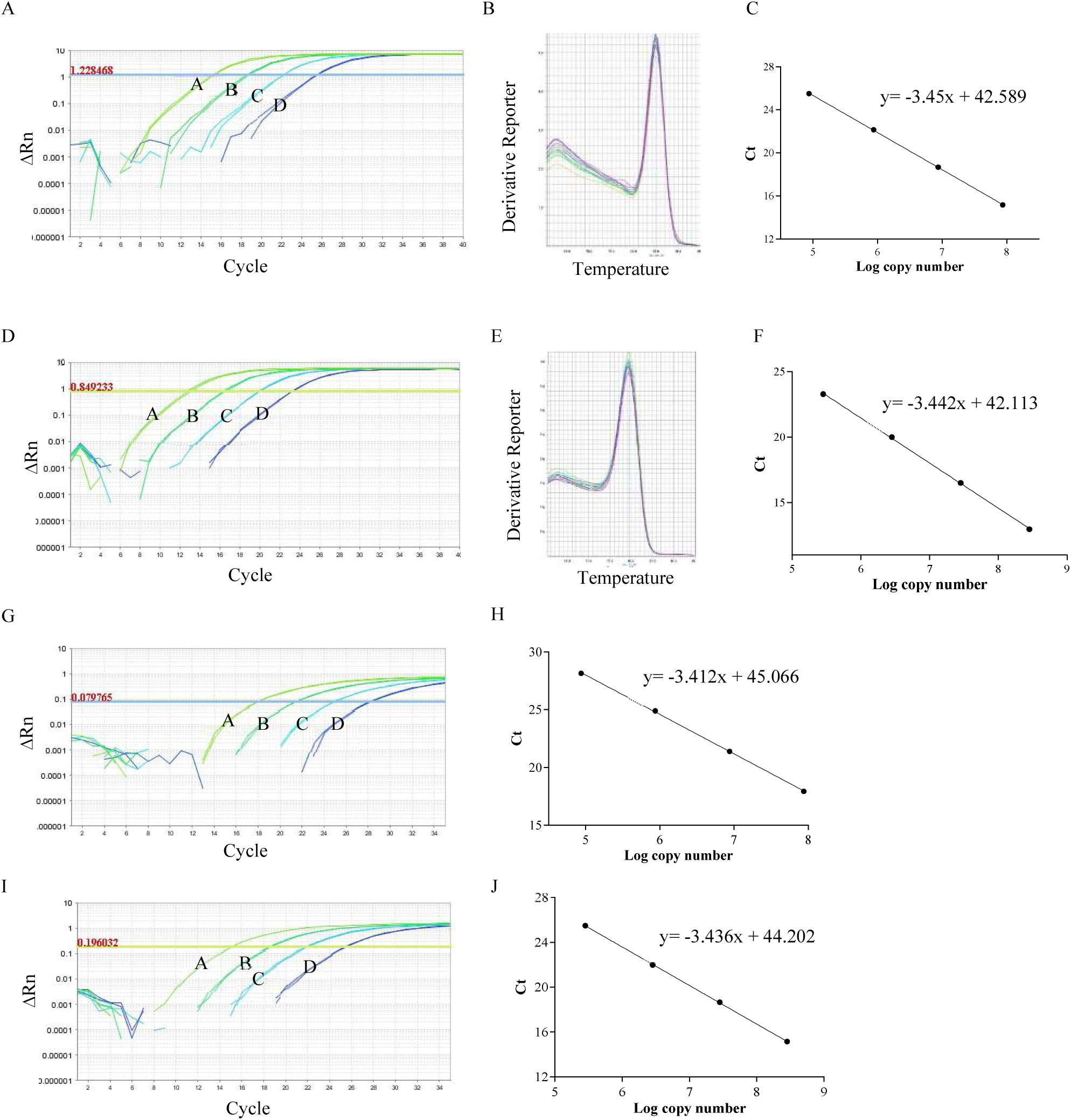
SYBR Green and probe based amplification (standards). **Panel A.** Amplification plot, GAPDH standard dilution: A: 8.69×10^7^, B: 8.69×10^6^, C: 8.69×10^5^, D: 8.69×10^4^. **Panel B.** Melt curve plot of standards **Panel C.** A standard curve created from panel A amplification plot demonstrates a strong linear regression between the Ct values and the DNA copy numbers (log scale). **Panel D.** Amplification plot, CAR standard dilution: A: 2.83×10^8^, B: 2.83×10^7^, C: 2.83×10^6^, D: 2.83×10^5^. **Panel E.** Melt curve plot of standards **Panel F.** A standard curve created from panel D amplification plot demonstrates a strong linear regression between the Ct values and the DNA copy numbers (log scale). **Panel G.** Amplification plot, GAPDH standard dilution: A: 8.69×10^7^, B: 8.69×10^6^, C: 8.69×10^5^, D: 8.69×10^4^. **Panel H.** A standard curve created from panel G amplification plot demonstrates a strong linear regression between the Ct values and the DNA copy numbers (log scale). **Panel I.** Amplification plot, CAR standard dilution: A: 2.83×10^8^, B: 2.83×10^7^, C: 2.83×10^6^, D: 2.83×10^5^. **Panel J.** A standard curve created from panel I amplification plot demonstrates a strong linear regression between the Ct values and the DNA copy numbers (log scale).

**FIGURE 3:**
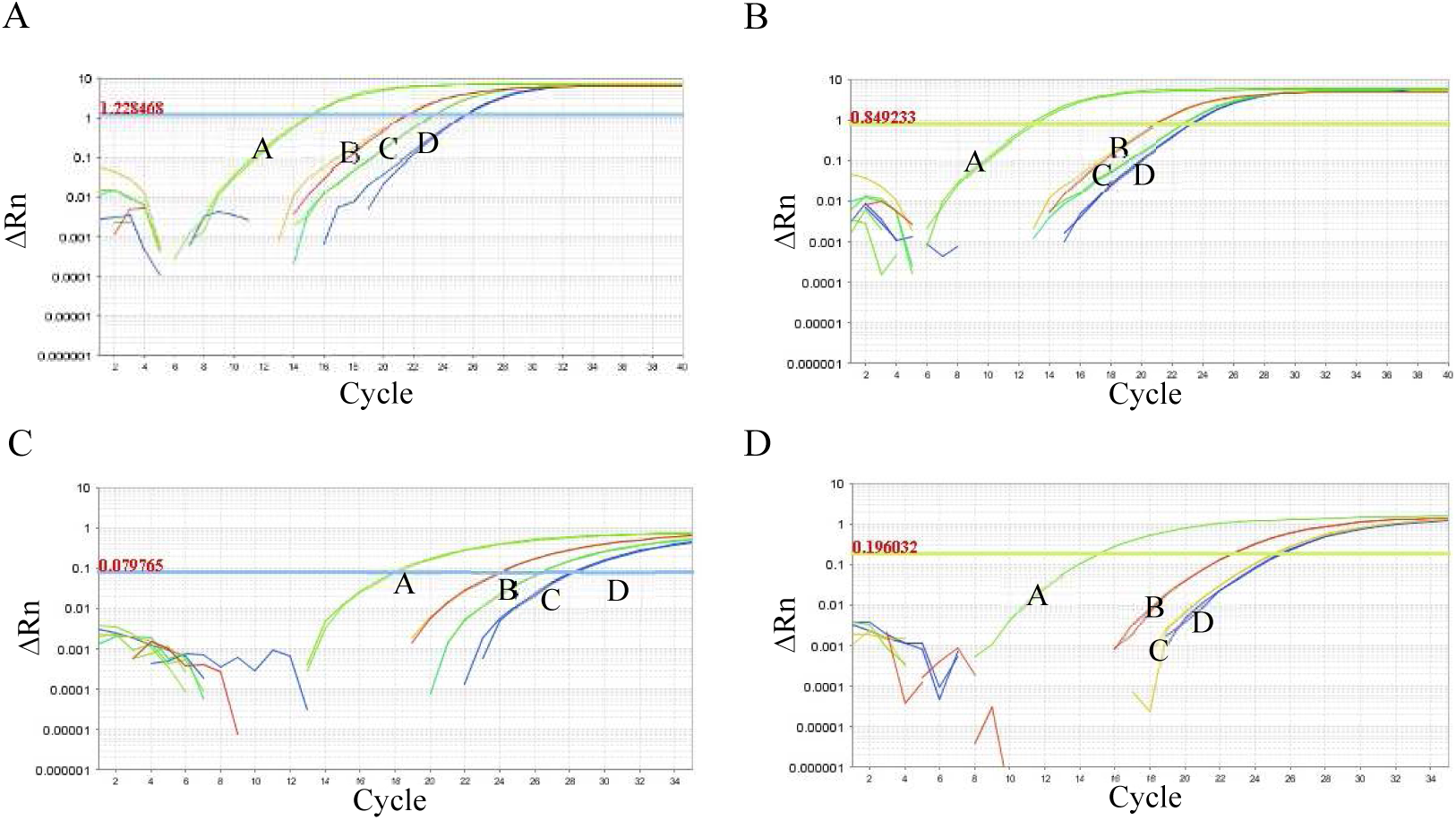
Amplification plot (standards and samples). **Panel A:** Amplification plot based SYBR Green, GAPDH standard and samples. A: 8.69×10^7^, B: CAR T cell 1∕2, C: CAR T cell 1∕10, D: 8.69×10^4^. **Panel B.** Amplification plot based SYBR Green, CAR standard and samples: A: 2.83×10^8^, B: CAR T cell 1∕2, C: CAR T cell 1∕10, D: 2.83×10^5^. **Panel C.** Amplification plot based probe, GAPDH standard and samples: A: 8.69×10^7^, B: CAR T cell 1∕2, C: CAR T cell 1∕10, D: 8.69×10^4^. **Panel D.** Amplification plot based probe, CAR standard and samples: A: 2.83×10^8^, B: CAR T cell 1∕2, C: CAR T cell 1∕10, D: 2.83×10^5^.

### Probe method

The amplification plot for GAPDH and CAR standards is shown in Figure 2G and I. The GAPDH standard curve demonstrates a strong linear regression with R_2_ = 0.9997, a PCR efficiency of 96.37%, and a regression *Pv* of 0.0001 (Figure 2H). The detected copy number in the GAPDH standard = 10^(Ct-45.066)/-3.412^. The detected copy number in a 1∕2 dilution of CAR T cells was calculated at 1.60×10^6^, and for a 1∕10 dilution it was 3.28×10^5^. The CAR standard curve also demonstrates a strong linear regression with R_2_ = 0.9999, a PCR efficiency of 95.45%, and a regression *Pv* < 0.0001 (Figure 2J). The detected copy number in the CAR standard = 10^(Ct-44.202)/-3.436^. The detected copy number in 1∕2 dilution of CAR T cell calculated 1.77×10^6^, and for the 1∕10 dilution was 3.61×10^5^. The copy number ratios of CAR T cells in 1∕2 and 1∕10 were 1.11 and 1.10 in total cells and 1.33 and 1.32 in CAR^+^ cells (83.6%), respectively (Table 2 and Figure 6 A-B). The amplification plot of CAR T cell samples alongside the first and last standard dilutions is shown in Figure 3 C-D. For copy number calculation based on the Probe method, two different dilutions of samples revealed approximately the same result; however, it seems more accurate than the SYBR Green method. The detected copy number for CAR and GAPDH standards also revealed precise calculations.

### Molecular viral titration

#### SYBR Green method

The amplification plot for CAR standards is shown in Figure 4A. The CAR standard curve also demonstrates a strong linear regression with R_2_ = 0.9967, a PCR efficiency of 103.53%, and a regression *Pv* < 0.0001 (Figure 4C). The detected copy number in a 1∕20 dilution of the viral sample was calculated at 5.43×10^5^, and for a 1∕50 dilution, it was 6.33×10^4^. The original concentration (copy/ml) was calculated at 2.90×10^9^ and 8.44×10^8^ for 1∕20 and 1∕50, respectively (Table 3). The amplification plot of viral samples alongside the first and last standard dilutions is shown in Figure 4D. For the SYBR Green method in viral titration, two different dilutions of the virus resulted in different viral titers, and based on the melt curve, any contamination reduced the accuracy of PCR (Figure 4B and E).

**FIGURE 4:**
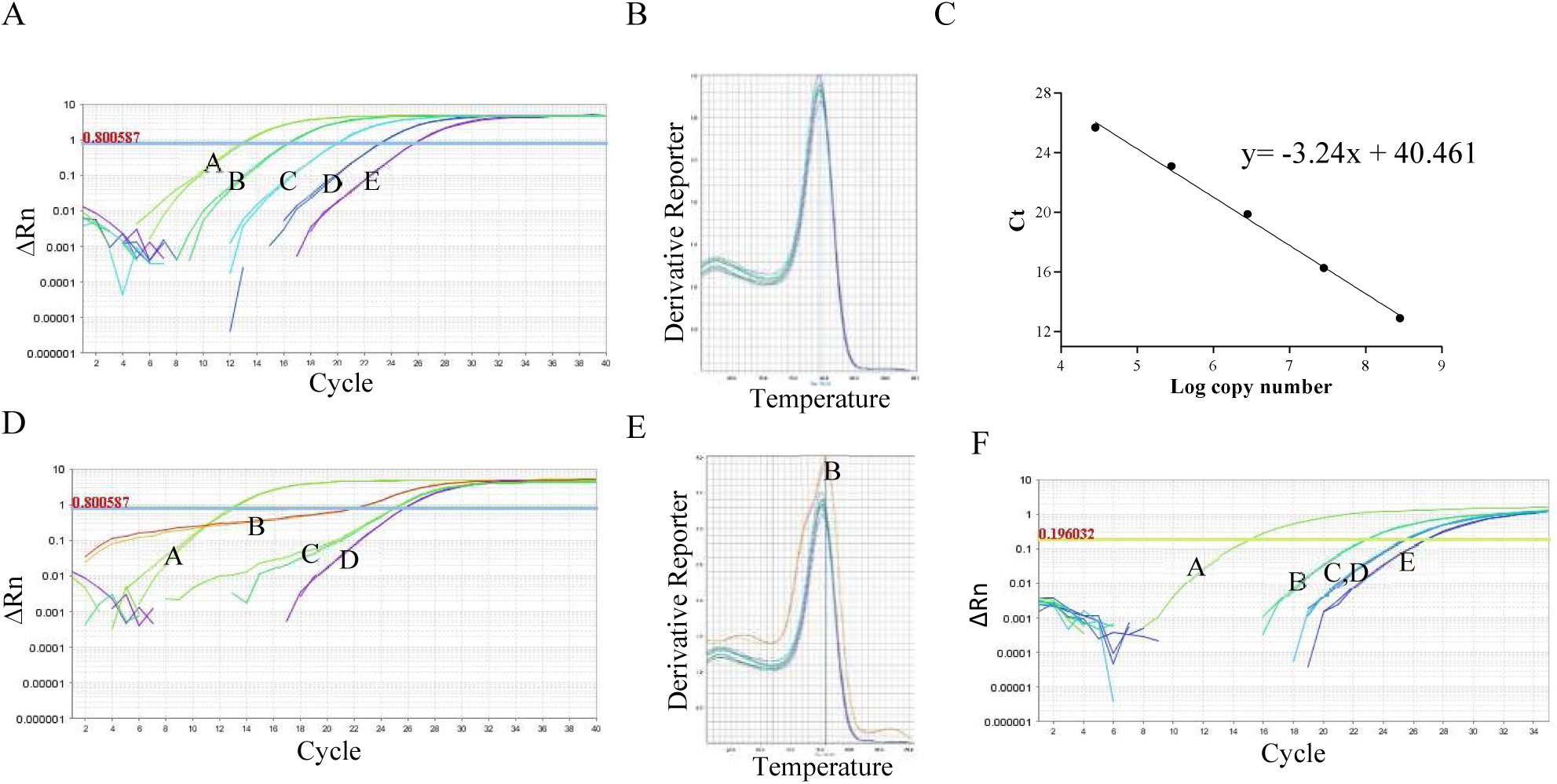
SYBR Green and probe based amplification (viral titration). **Panel A:** Amplification plot, CAR standard dilution: 2.83×10^8^, B: 2.83×10^7^, C: 2.83×10^6^, D: 2.83×10^5^, E: 2.83×10^4^. **Panel B.** Melt curve plot of standards and samples. **Panel C.** A standard curve created from panel A amplification plot demonstrates a strong linear regression between the Ct values and the DNA copy numbers (log scale). **Panel D.** Amplification plot based SYBR Green, CAR standard and samples: A: 2.83×10^8^, B: virus 1∕20, C: virus 1∕50, D: 2.83×10^4^. **Panel E.** Melt curve plot of standards and samples, B: virus 1∕20. **Panel F.** Amplification plot based probe, CAR standard dilution: A: 2.83×10^8^, B: virus 1∕2, C virus 1∕20, D: 2.83×10^5^, E: virus 1∕50.

**Table 3:**
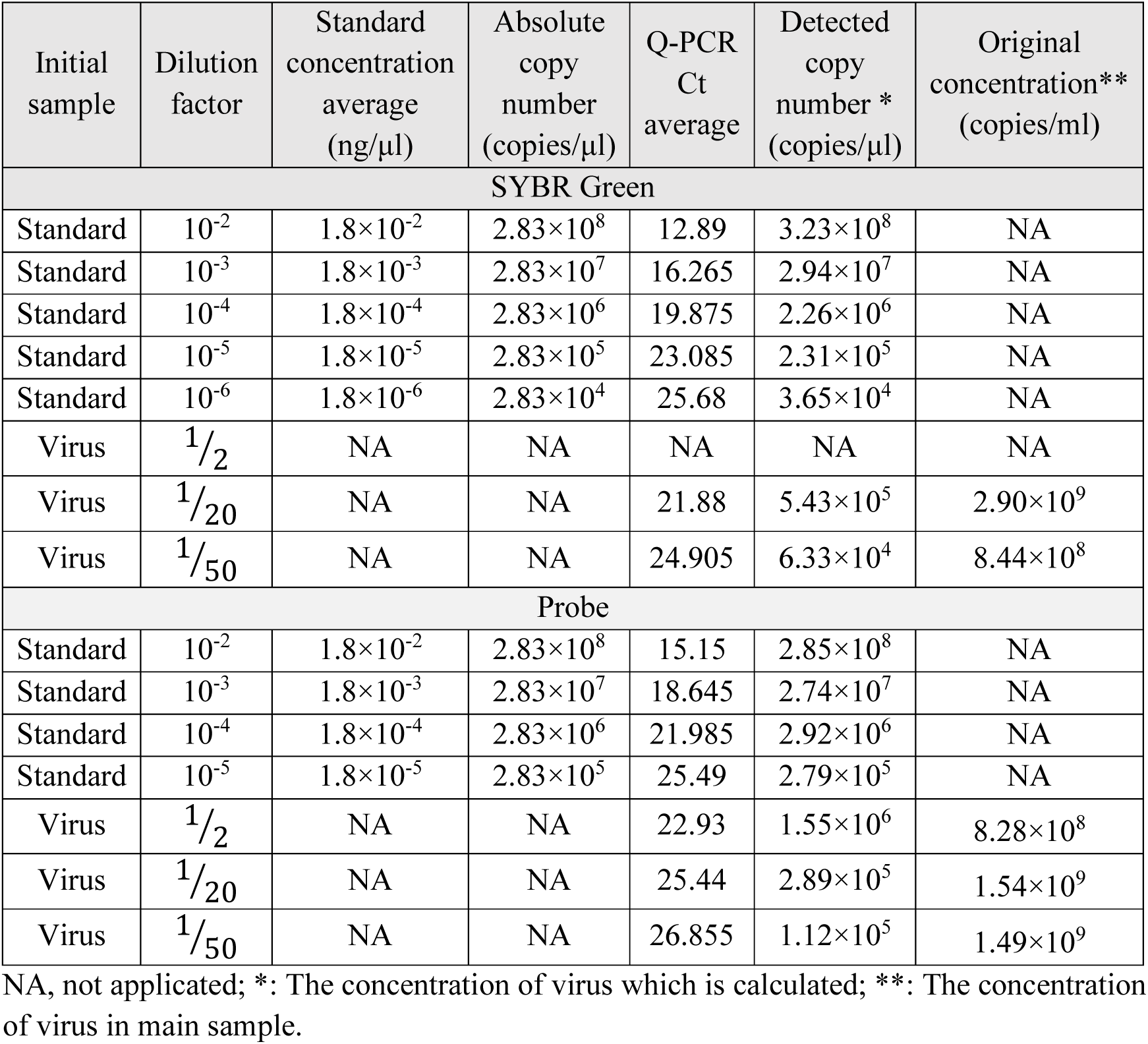
Viral titration SYBR Green and Probe (sample1)

#### Probe method

The amplification plot of CAR standards for Probes was the same for Probe based copy number detection. The detected copy numbers in 1∕2, 1∕20, and 1∕50 were 1.55×10^6^, 2.89×10^5^, and 1.12×10^5^, respectively. The original concentration (copy/ml) was calculated at 8.28×10^8^, 1.54×10^9^, and 1.49×10^9^ for 1∕2, 1∕20, and 1∕50, respectively (Table 3). The amplification plot of viral samples alongside the first and last standard dilutions is shown in Figure 4F. For viral titration based on the Probe method, three different dilutions of samples revealed approximately the same result, and the same contamination in samples had no adverse effect on PCR efficiency with Probe based methods.

#### SYBR Green vs. Probe methods difference

The sample contamination could significantly reduce the accuracy of viral titration tests whit more dependency on SYBR Green methods but no significant difference was reported in VCNs with the two methods (Table 5, Figure 5 A-B).

**FIGURE 5:**
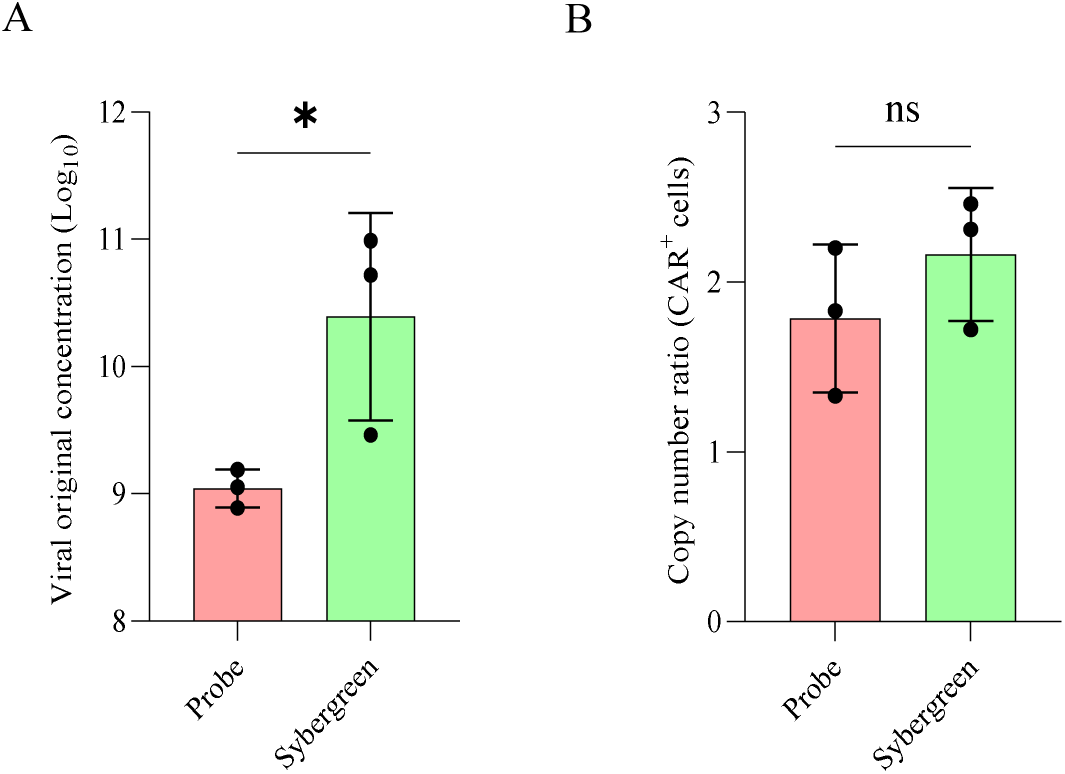
SYBR Green vs. probe. **Panel A.** Viral original concentration (log10-copies/ml) in different methods were evaluated. There is significant difference between two methods. **Panel B.** Copy number ratio (CAR copies/cell) in different methods were evaluated. There is no significant difference between two methods. The unpaired t-test was assessed between groups.

#### Cellular viral titration

In the analysis of viral samples, the 1∕20 and 1∕40 dilutions of samples resulted in >99% T cells (CD3^+^), 13.3% CAR T cells for the 1∕20 sample dilution, and 5.46% CAR T cells for the 1∕40 sample dilution. Original virus titers were 1.73 × 10^6^ (TU/ml) and 2.13 × 10^6^ (TU/ml) for 1∕20 and 1∕40 dilutions of samples, respectively (Figure 6 and Table 4). Viral titration based on cellular transduction revealed functional viral titration; however, molecular viral titration resulted in only genome detection, which means lower viral titration in cell-based strategies, consistent with our results.

**FIGURE 6:**
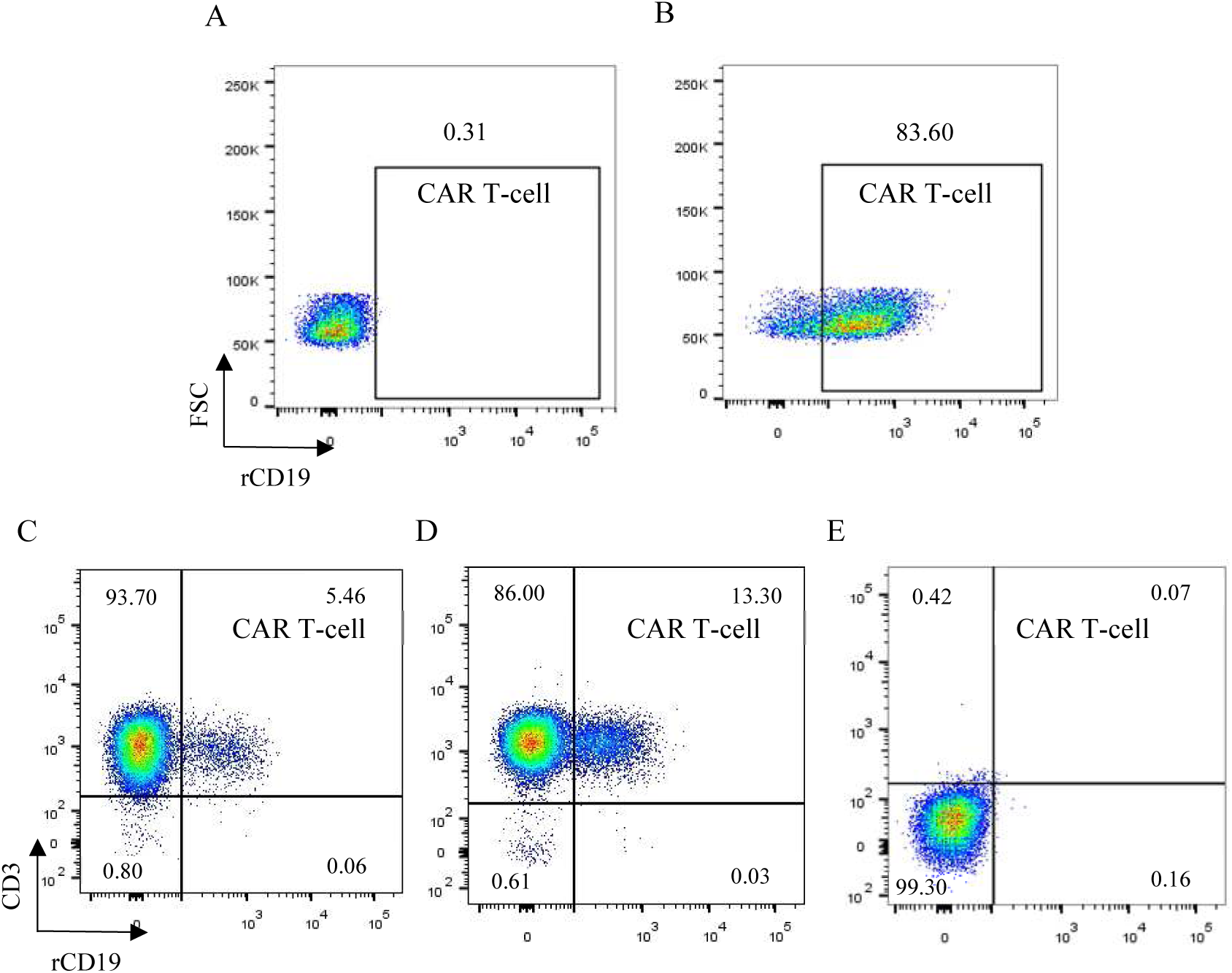
Viral transduction and titration. **Panel A.** Iso type control. **Panel B.** virus transduction. 83.6% CAR T cell. **Panel C.** 1∕20 dilution of virus. 5.46% CAR T cell. **Panel D.** 1∕10 dilution of virus, 13.3% CAR T cell. **Panel E.** Iso type control. rCD19: recombinant CD19.

**Table 4:** Cellular viral titration (sample1)

| Dilution | 0.025 | 0.05 |
|---|---|---|
| DF | 40 | 20 |
| F | 0.054% | 0.133% |
| Cn | 8 x 10<sup>4</sup> | 8 x 10<sup>4</sup> |
| V (ml) | 0.1 | 0.1 |
| Titer (TU/ml) | 1.73×10<sup>6</sup> | 2.13×10<sup>6</sup> |

DF: The virus dilution factor; F: The frequency of GFP-positive cells determined by flow cytometry, the range between 1 to 20 were acceptable; Cn: The total number of target cells infected; V: The volume of the inoculum.

**Table 5:** Results summary (sample1-3)

| Initial sample | Original concentration* (copies/ml) | Original concentration* (copies/ml) | Titer (TU/ml) | Copy number ratio** (CAR<sup>+</sup> cells) | Copy number ratio** (CAR<sup>+</sup> cells) |
| Methods | SYBR Green | Probe | Cellular titration | SYBR Green | Probe |
|---|---|---|---|---|---|
| Virus 1 | 2.90×10<sup>9</sup> | 1.54×10<sup>9</sup> | 1.73×10<sup>6</sup> | NA | NA |
| Virus 2 | 5.24×10<sup>10</sup> | 1.13×10<sup>9</sup> | 2.16×10<sup>6</sup> | NA | NA |
| Virus 3 | 9.76×10<sup>10</sup> | 7.73×10<sup>8</sup> | 2.49×10<sup>6</sup> | NA | NA |
| CAR T cell 1 | NA | NA | NA | 1.72 | 1.33 |
| CAR T cell 2 | NA | NA | NA | 2.31 | 1.83 |
| CAR T cell 3 | NA | NA | NA | 2.46 | 2.20 |

NA, not applicated; \*: The concentration of virus in main sample; \*\* The mean copy number of CAR in each cells CAR/GAPDH in CAR<sup>+</sup> cells.

## Discussion

CAR T cells targeting the CD19 antigen are currently being developed in clinical trials and revolutionizing the treatment of patients with hematological disorders. These CAR T cells, approved by the FDA for clinical use, typically use viral vectors, primarily γ-retrovirus and lentivirus (17). These viruses belong to the Orthoretrovirinae subfamily of Retroviridae, converting RNA into cDNA that integrates randomly into the cellular genome. This integration allows for long-term expression of the transgene in target cells (18, 19). Nonetheless, the utilization of viral vectors in genetically modified organism (GMO) products, exemplified by CAR T cells, for therapeutic applications mandates stringent biosafety quality control (QC) assessments (6). Genotoxicity testing using various methods is essential to confirm that the random integration of viral cDNA does not harm cell cycle-related genes, potentially leading to tumor formation post-infusion (20). Additionally, it is important to determine the average number of viral cDNA copies integrated (VCN) in CAR T cell products (7). An increased VCN may be associated with greater genetic integration and a higher risk of genotoxicity (14). The FDA recently highlighted the serious risk of T cell malignancies associated with all currently approved CAR T cell therapies. Another critical QC measure involves viral vector titration and MOI, both of which impact VCN. Increasing the MOI leads to higher VCN, necessitating careful characterization and limitation within a specific range during CAR T cell manufacturing (14).

Here, we developed a Q-PCR assay based on ACM for both VCN and viral titration. Q-PCR is a widely used tool in laboratories and is easily accessible, with many professionals skilled in PCR-related techniques. However, there are some challenges with this method that are related to standards and require attention to practical and technical issues. The creation of standard samples containing a CAR T cell clone with a defined and stable VCN of CAR highlighted some important considerations in Q-PCR methods. We used any CAR T cell DNA for the generation of both CAR and GAPDH standards, as they were readily available. The same primers and standards were used for both VCN and viral titration experiments, allowing for examination under the same PCR conditions. The primers were designed in U5 LTR and could be utilized not only for evaluating CAR T cell copy numbers or CAR viral titration, but also for other target cells or viruses with the same SFG backbone. There are two common methods in Q-PCR: SYBR Green and Probe based methods. While SYBR Green is more cost-effective, we also analyzed the same samples with a Probe to compare the two methods.

In this experiment, CAR T cell DNA was used to produce GAPDH and CAR standard. The standard linear regression was highly strong, with R_2_ > 0.99, PCR efficiency ranging from 94.91% to 103.53%, and *Pv* ≤ 0.0003 for the GAPDH and CAR standards in both SYBR Green and Probe based experiments. Based on the standard results, both Probe and SYBR Green methods were reliable for VCN and viral titration experiments. For VCN calculation, the SYBR Green method resulted in 1.72 and 1.67 copy numbers in two dilutions, which were somewhat similar. Similarly, the Probe based VCN calculation showed results of 1.33 and 1.32 in two dilutions, with the Probe based calculations yielding more precise results. When conducting viral titration using SYBR Green method, different results were obtained for the two dilutions. Melt curve analysis showed that the 1/20 dilution had an improper curve, while the 1/50 dilution yielded better results. In Probe based titration, all dilutions showed similar results, except for the 1/2 dilution which had a slight difference. It appears that the purity of samples was crucial to the final results, with the Probe based strategy showing significantly better results, and having lower dependency on sample contamination. Therefore, we recommend using Probe based methods with different diluted samples to compare final results for data reporting. Viral titration experiments could also be analyzed using cellular methods. In this experiment, the same viral samples resulted in titrations differing by around 100-1000 folds. Molecular-based methods analyze any DNA copies, while cell-based methods only analyze functional virus. The variability in transduction methods, such as Retronectin or Polybrene, led to different transduction ratios, with differences at any stage of transduction resulting in varying ratios. In analyzing transduction using T cells, it is important to note that for CAR T cell production in clinical settings, the same cell line should be used to analyze different viral batches. Variations between cell lines can also lead to different transduction ratios. The stability of the virus during incubation at 37°C until gathering and freezing can impact the functionality of the virus for cell-based viral titrations. It is essential for molecular-based methods characterizing viral batches to show consistency across different experiments.

## Conclusion

In this study, the VCN and viral titration analyses were developed using the same primers targeting the U5 region in the 5’ LTR. These primers could also be utilized for other transduced cells and various retroviral samples with the SFG backbone. Both SYBR Green and Probe based Q-PCR were employed, indicating that the Probe strategy exhibited less susceptibility to sample contamination and closer results to cellular based titration for viral titration.

## Author Contributions

Hamidreza Ebrahimiyan: Investigation, methodology, formal analysis, data curation and writing original draft. Ali Sayadmanesh: writing – review and editing and data curation. Mohsen Basiri: Conceptualization, writing – review and editing, data curation and supervision. Hossein Baharvand: funding acquisition, review, editing and supervision.

## Acknowledgement

We deeply acknowledge all donors who supported this study. We also thank the staff and technicians at the Royan Institute for Stem Cell Biology and Technology for their technical assistance.

## Data Availability Statement

The data generated and analyzed during the study are available as requested.

## Declaration of generative AI and AI-assisted technologies in the writing process

During the preparation of this work the authors used ChatGPT (OpenAI) in order to correct spelling and grammar errors and improve readability of the text. After using this tool, the authors reviewed and edited the content as needed and take full responsibility for the content of the published article.

